# Inferring transmission bottleneck size from viral sequence data using a novel haplotype reconstruction method

**DOI:** 10.1101/2020.01.03.891242

**Authors:** Mahan Ghafari, Casper K. Lumby, Daniel B. Weissman, Christopher J. R. Illingworth

## Abstract

The transmission bottleneck is defined as the number of viral particles transmitted from one host to another. Genome sequence data has been used to evaluate the size of the transmission bottleneck between humans infected with the influenza virus, however, the methods used to make these estimates have some limitations. Specifically, approaches using viral allele frequency data may not fully capture a process which involves the transmission of entire viral genomes. Here we set out a novel approach for inferring viral transmission bottlenecks; our method combines haplotype reconstruction, a method for inferring the composition of genomes in a viral population, with two maximum likelihood methods for bottleneck inference, tailored for small and large bottleneck sizes respectively. Our method allows for rapid calculation, and performs well when applied to data from simulated transmission events, being robust to errors in the haplotype reconstruction process. Applied to data from a previous household transmission study of influenza A infection we confirm the result that the majority of transmission events involve a small number of viruses, albeit with slightly looser bottlenecks being inferred, with between 1 and 13 particles transmitted in the majority of cases. While influenza A transmission involves a tight population bottleneck, the bottleneck is not so tight as to universally prevent the transmission of within-host viral diversity.

## Introduction

Viral populations experience large fluctuations in population size. During the course of an infection many thousands of viruses may be produced by each infected cell (1), yet in the process of transmission only a small number of viruses may get through to found a new infection (2). The size of the bottleneck undergone by a viral population at the moment of transmission has an important impact on the evolution of that virus. Where larger numbers of viral particles are involved in transmission, a greater amount of genetic diversity is preserved between hosts; where smaller numbers of particles are transmitted, between-host evolution becomes more of a stochastic process (3). Studying transmission at the scale of individual hosts therefore gives an insight into larger-scale patterns of viral evolution.

Genetic data provides an invaluable insight into processes of viral evolution (4). Such data has been at the core of a variety of approaches for the quantitative analysis of population bottlenecks, typically using observations of minority variants, or their allele frequencies, to make a statistical inference. For example, counting the number of minority variants shared between hosts can be informative of whether transmission occurred between specific hosts (5, 6). If the route of transmission is known, the same count can be used to estimate the size of the population bottleneck (7). A model of genetic drift may also be applied: smaller or larger changes in the composition of a viral population suggest that a larger or smaller number of viruses were transmitted (3, 8–11). In some situations, engineered viruses with genetic markers have been used to directly evaluate transmission events (12, 13)

Recent studies of influenza transmission between human hosts have used metrics based upon changes in allele frequencies to evaluate the bottleneck at transmission (3, 11, 14, 15). Such metrics have limitations; transmission is ultimately an event in which whole viruses, rather than independent alleles, are passed from one host to another. Neglecting genetic linkage in this way can skew the results of inference methods (16). Inspired by this, a recent study on the assessment of viral transmissibility used sequence data to evaluate transmission at the level of viral genomes (17).

Accounting for genetic linkage between alleles becomes more difficult as the diversity of a viral population increases. In modelling the action of selection on a diverse population, the large number of potential genome sequences can make calculations infeasible. Considering cases in which selection among transmitted variants is not the dominant effect at transmission (3) we here set out an alternative approach for the inference of population bottlenecks, incorporating the true genetic structure of viruses. Our approach has two components. Firstly, given sequence data collected before and after a transmission bottleneck, we apply a method of haplotype reconstruction, using a maximum likelihood framework to calculate a parsimonious reconstruction of the viral population, as observed before and after transmission.

A broad variety of computational tools have previously been described for the purpose of haplotype reconstruction in various contexts (18–23); ours fits naturally into the bioinformatic framework we have outlined in previous publications (24, 25). Secondly we use the haplotype reconstruction to infer a bottleneck size at transmission; our framework contains two alternative approaches optimised for smaller and larger bottleneck sizes respectively. We test our method against simulated data describing viral transmission events with a broad range of population bottlenecks. Finally, we re-evaluate data from a previous study of influenza transmission between human hosts (3). Our study supports the hypothesis of a generally small transmission bottleneck for influenza viral populations (3, 26) albeit with fractionally higher bottleneck sizes inferred from the same data.

## Results

Before considering data from a study of human infection, we applied our method to simulated transmission data, examining the haplotype reconstruction and bottleneck inference steps.

### Allele-based versus haplotype-based inference

An initial example of a transmission bottleneck highlights the potential pitfalls of the use of single-allele models for evaluating transmission (Figure 1). In this simulated system data were collected from before and after a transmission bottleneck. While during transmission the viral population changed substantially at the genotype level, these changes were not fully reflected in the allele frequency data from each population. As a consequence, inferences of the bottleneck at transmission, calculated using haplotype- and allele-frequency methods, differed by close to two orders of magnitude. Although the population used in this analysis is something of an artificial construction, the result highlights a fundamental point of biology. Rather than independent alleles, viral transmission involves the transmission of complete viral genomes, and approaches which neglect this may be flawed in their outcome. We are therefore motivated to consider the transmission of viruses on the genotype level.

**Fig. 1.**
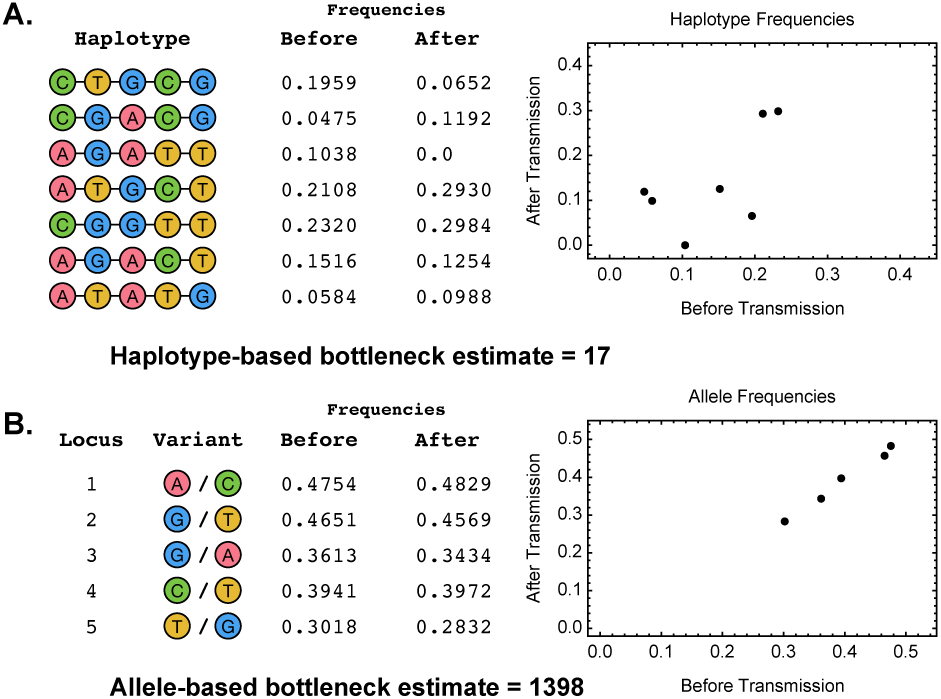
**A.** Simulated system of viral transmission. A population comprising seven viral genotypes transmits to a new host, leading to a population in the recipient which includes six of the seven genotypes. A plot shows the sampled frequencies of the distinct genotypes, or haplotypes, before and after transmission, reported to four significant figures. Our explicit model of viral transmission based on haplotype frequencies (described in the text) infers a population bottleneck of 17 viruses from these data. **B.** An alternative analysis of the same population samples allele frequencies from the population before and after the transmission event; these are shown in an equivalent plot. A calculation of the population bottleneck from these data infers a value nearly two orders of magnitude larger than that of our previous calculation.

### Haplotype reconstruction

Applied to simulated data our method showed a reasonably good ability to correctly identify haplotypes with a correct inference of all haplotypes being made in more than half of the cases tested (Figure 2). Our approach uses a maximum likelihood method to infer the most parsimonious reconstruction of a viral population, given sequence data. To test our approach we simulated data describing the transmission of an influenza viral population, from a host to a recipient individual. Each segment in the population was modelled as containing six distinct haplotypes, applying a method for generating data described in a previous study (17). Simulated sequence data from the viral populations in each host were used to infer which haplotypes were present in the transmission event and their frequencies. Combining data across segments the most common outcome was a correct reconstruction of all of the haplotypes in the population.

**Fig. 2.**
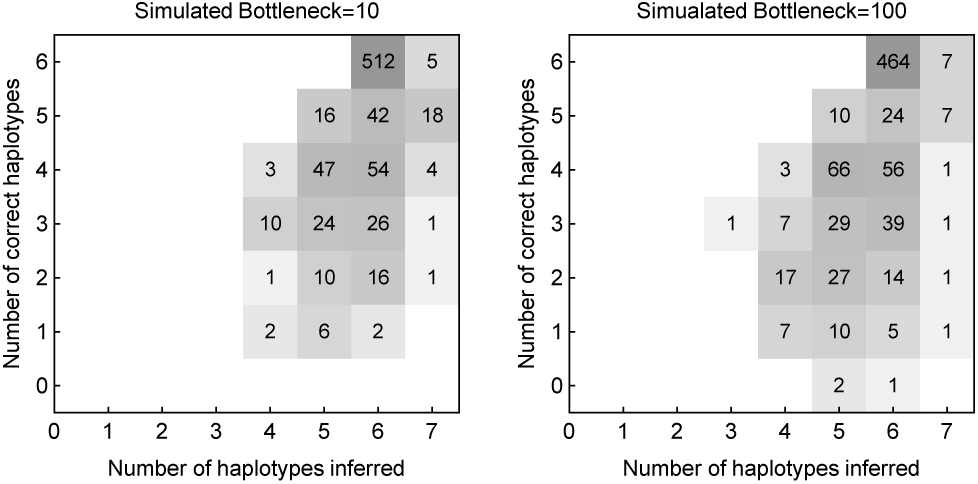
Numbers of inferred and correctly inferred haplotypes given simulated sequence data. A total of 6 haplotypes were included in each of 800 simulations tested.

### Inference of population bottlenecks

Our two methods for bottleneck inference produced good results when applied to simulated viral transmission data (Figure 3). As described in the Methods section, the two methods we apply generalise in turn the approaches of two previously-described single-locus methods for bottleneck inference (11, 14). Our “compound method” uses a model of genetic drift in a continuous space of genotype frequencies, in which smaller changes in frequencies correspond to a lesser amount of stochasticity in transmission, and hence a larger population bottleneck (14). Our “explicit method” explicitly evaluates all of the possible outcomes of a transmission event across a discrete space: the fact that an integer number of viruses of each genotype are transmitted is used to weigh up the likelihood of different potential bottlenecks (11).

**Fig. 3.**
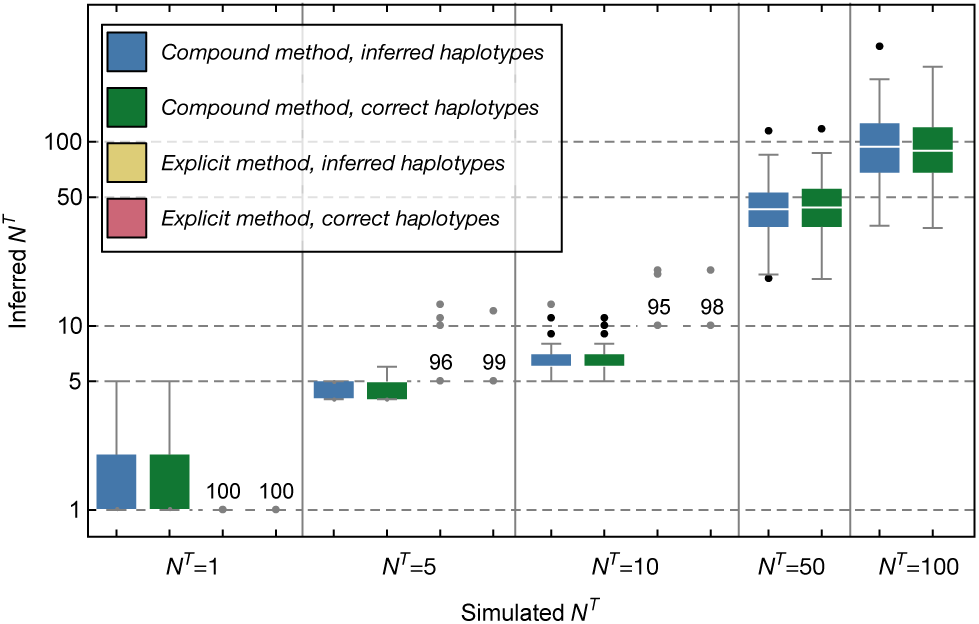
Transmission bottleneck sizes inferred from simulated data using different input data and methodologies. Inferences are shown in colour according to the data and method used. Calculations with inferred haplotypes took as input data generated from a haplotype reconstruction method applied to simulated sequence data, in which both the haplotypes and their frequencies before and after transmission were inferred. Calculations with the correct haplotypes took as input data from a haplotype reconstruction in which the identities of the correct haplotypes were given, with only their frequencies being inferred. Inferences from the explicit method were only calculated for smaller population bottleneck sizes as the method does not scale well to evaluating larger bottlenecks. Results from the explicit method were so accurate as to not have a meaningful interquartile range: numbers displayed in these cases indicate the number of inferences giving a precisely-correct inference of the population bottleneck. Horizontal dashed lines indicate the simulated bottleneck sizes.

Applying these methods to simulated data, the compound method generally did well, inferring transmission bottlenecks that were close to the simulated values. One advantage of this method is that its running time does not increase with the bottleneck size, enabling the analysis of very high potential bottleneck sizes. A disadvantage of the method is that, despite improvements made with respect to its predecessor (17), the mathematical approximations made in its construction mean that it does not always perform so well at low bottleneck sizes, producing a visible underestimate of bottlenecks of size 10. Further inferences of bottleneck size were made using reconstructions of haplotypes in which the correct simulated haplotypes were pre-specified, learning only their frequencies. Using these improved data did not produce a noticeable improvement in the inference of the bottleneck size, suggesting that our inference of bottleneck size is robust to errors that arise from our haplotype reconstruction method. Bottleneck sizes in each case were calculated across eight independent viral segments.

Given our simulated data, the explicit method outperformed the compound method at low bottleneck sizes, inferring exactly correct values in the majority of cases with very little error. A disadvantage of the explicit method is that in requiring the evaluation of all possible outcomes of a transmission event, the computational time it requires grows very rapidly as the bottleneck size increases. For this reason, we did not apply it to data from higher simulated population bottlenecks. As with the compound method performance did not greatly improve given frequencies inferred using the correct viral haplotypes; errors in haplotype reconstruction did not have a strong effect on the inferred bottleneck sizes.

Inference of bottleneck size for a segment was not possible in two cases. Firstly, if our haplotype reconstruction found evidence for only a single viral haplotype, no inference was possible, insufficient information about the event being available. Secondly, if the viral population in the recipient was inferred to have arisen purely from a *de novo* haplotype, which had swept to fixation in the population between the establishment of the infection and the collection of the sequence data, this result was uninformative in identifying a bottleneck. In either of these circumstances, data from a viral segment was ignored, inferences conducted for the remaining segments being combined to infer the final bottleneck size.

In considering the differences in inferences achieved by the two methods at low bottleneck sizes, it is perhaps helpful to consider the simple case where a single allele frequency changes from 50% frequency in the donor to 5% in the recipient. Within the compound method this represents a large change in allele frequency, corresponding to a large amount of genetic drift, and hence a low bottleneck size. By contrast under the explicit method variation at a frequency of 5% is difficult to achieve given a low population bottleneck; a case in which 20 viruses were transmitted, one of which had the minority variant, would give a more coherent explanation

### Application to data from a household study

Our transmission model was applied to data collected from a previously published household study (3). This study used a single-locus inference model to identify narrow bottlenecks in human-to-human transmission, with all but a single event being inferred to involve the transmission of between one and four viral particles. Short-read data from this study was filtered and processed into variant data before being fed into our method. Having identified polymorphic loci in pairs of transmission data using an allele frequency cutoff of 2% we generated multi-locus reads from the data using the SAMFIRE sofware package (25), using these to generate an inference of haplotype frequencies before and after transmission. These frequencies were used to infer population bottleneck sizes for each transmission event.

We confirm the previous inference of tight population bottlenecks in all cases (Figure 4). In the majority of transmission events (29 out of 38 events for which we obtained an inference), bottlenecks of size *N*^*T*^ = 1 were inferred by both of our methods, consistent with all of the diversity of the viral population in the original host being lost at transmission. While not implying that these infections were started by a single viral particle, these results are consistent with the hypothesis of a generally tight bottleneck at transmission. In eight out of the remaining nine transmission events, intermediate bottleneck sizes were inferred, with a range from 2 to 7 in the compound method and from 2 to 13 in the explicit method. Evidence from simulated data suggests that the explicit method is probably more accurate in this range. Finally, there was a single case in which a bottleneck size of 200 or more was inferred; this was set as the upper limit considered by our study. Our inference in this case matched the original analysis of the data. A further statistical analysis of the samples collected before and after transmission indicated a greater degree of similarity between allele frequencies than was previously found in a case where replicate clinical samples were processed and sequenced in parallel (27). Whereas in the previous study, measurements of allele frequencies from samples split from the cDNA synthesis step onwards were consistent with an effective read depth (that is equivalent to an error-free sample depth) of one thousand or more, here an effective depth in excess of 20,000 was inferred, demonstrating that the before- and after-transmission samples were extremely similar. This case could represent either a very unusual transmission event, in which an extreme number of viruses were transmitted, or potentially an isolated error in the processing of a large number of sequence samples.

**Fig. 4.**
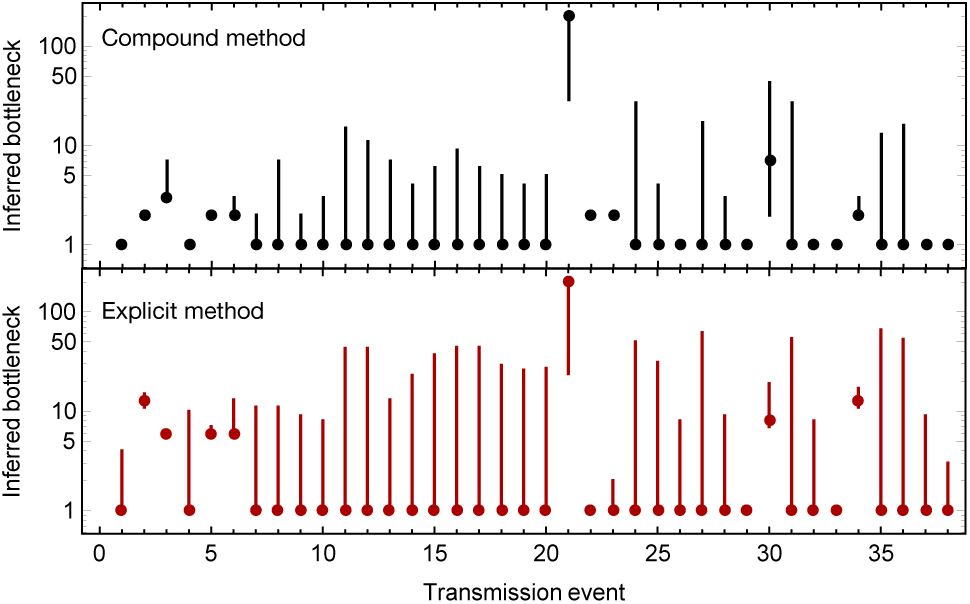
Bottleneck sizes inferred from the data presented by (3). Dots indicate the maximum likelihood bottleneck size inferred for each of the 38 systems in this work for which we were able to infer a bottleneck. Vertical bars represent confidence intervals equivalent to a cut-off of 2 log likelihood units.

Cases in which the explicit method inferred larger bottleneck sizes than the compound method could be explained in terms of the preservation of allele frequencies at relatively low frequencies; as explained above the explicit method can favour a higher bottleneck in such cases.

Our approach was not able to infer a population bottleneck in five of the transmission cases analysed by the original study. In these cases a low level of polymorphism observed before transmission was no longer present after transmission. Application of our haplotype reconstruction method in these cases did not find statistical evidence for more than one haplotype (plus noise) in these systems, at least two specific haplotypes being required for an inference of bottleneck size. We understand this in terms of our haplotype reconstruction method being less sensitive to detecting variation than is the 2% allele frequency cutoff used in the original study; the presence of a variant allele at 2% frequency was not always sufficient evidence for our code to infer the existence of two specific genetic variants in the population. In these cases, the loss of host genetic variance at transmission would lead our methods to the conclusion that a bottleneck of *N*^*T*^ = 1 best explained the observed data, strengthening our main result of a tight bottleneck size. The sensitivity of our method in calling additional haplotypes can be somewhat arbitrarily tuned.

## Discussion

We have here set out a haplotype-based approach for the inference of transmission bottlenecks, and demonstrated its application using data from a study of transmission of influenza A infection. Our approach uses a haplotype reconstruction approach to infer the composition of the viral population before and after transmission; by requiring substantial evidence to add an additional haplotype to the model, the approach limits the complexity of the inferred viral population, improving the feasibility of bottleneck inference relative to a previous approach (17). While our haplotype reconstruction method was not perfect in reproducing the details of a viral population, errors resulting from this method did not greatly harm our inference of population bottleneck sizes.

Our inference of bottleneck size comprises two distinct methods, optimal for distinct transmission bottleneck sizes. The first of these generalises the approach of Poon et al. (14), who used a formula based on genetic drift to evaluate changes in allele frequencies. Our compound method generalises this to changes in haplotype frequencies, which occur in higher dimensional sequence space; it further incorporates uncertainty in the inferred haplotype frequencies and genetic drift arising from within-host population growth. This method has the advantage of being rapid to calculate at high bottleneck sizes, but potentially underestimates bottleneck sizes at low values of *N*^*T*^. Our second method, the explicit method, generalises the approach of Sobel Leonard et al. (11), who apply a beta-binomal formula to evaluate possible discrete outcomes of a transmission process. In spirit we repeat this approach, summing a likelihood function over the set of possible outcomes of a transmission of viral haplotypes. This approach is limited in its application to systems of higher complexity, becoming slow where there are many haplotypes or where *N*^*T*^ is large, but is likely more accurate at lower bottleneck sizes.

Some challenges remain in the inference of population bottleneck sizes from within-host sequence data. In particular, the dynamics of the very early stages of population growth, from the initial founder viruses to the large population typical of influenza infection, are not necessarily well understood. Knowledge of the extent to which this affects the genetic composition of the viral population would improve the potential for accurate inference. We note that, where ethically feasible, the use of neutral markers provides a more direct approach for evaluating transmission events (12).

The biological conclusion of our study was that, applying an improved inference method to sequence data from a household study of influenza A infection, we largely replicate the finding that influenza A transmission involves a small number of viral particles, albeit that our results have a longer tail of bottleneck sizes inferred to be greater than 1; using our approach we obtained estimates that up to 13 viruses were transmitted. While transmission therefore limits the inheritance of viral diversity, its effect in doing so is not absolute; cases of the transmission of viral diversity do exist and may have some influence on broader viral evolutionary dynamics.

## Methods

### Notation

A guide to the notation used in our methods is shown in Figure 5. Briefly, we represent the populations before and after transmission by vectors of unknown haplotype frequencies, referred to as ***q***^*B*^ and ***q***^*A*^ respectively. These are separated by transmission with a bottleneck *N*^*T*^, forming the founder viral population ***q***^*F*^ in the recipient, then within-host growth, represented in our model by a single generation of genetic drift with effective size *N*^*G*^. The unknown vectors ***q***^*B*^ and ***q***^*A*^ are indirectly observed via the datasets ***x***^*B*^ and ***x***^*A*^, which are used to generate the estimated haplotype frequencies ***q***\*^*B*^ and ***q***\*^*A*^.

**Fig. 5.**
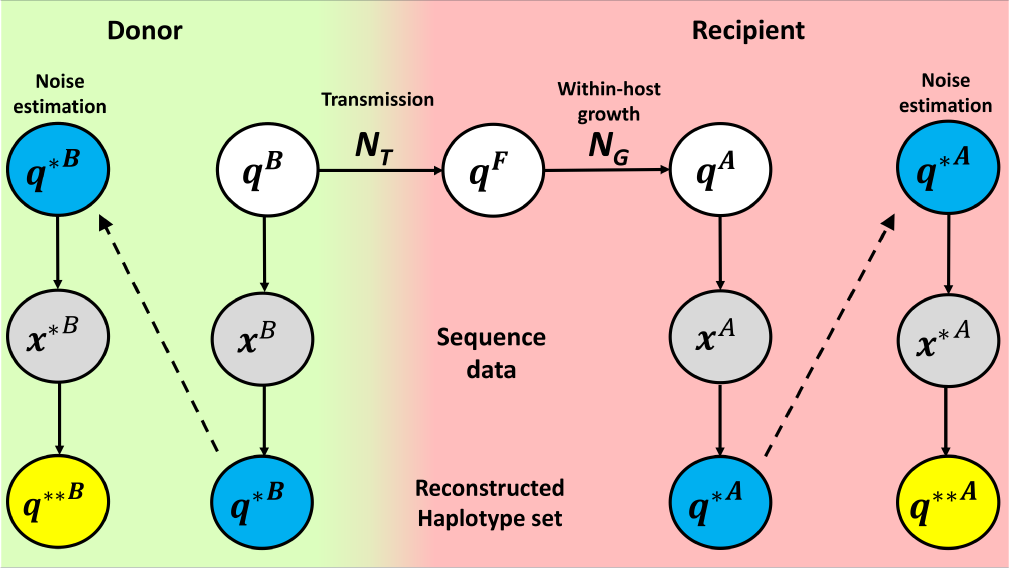
Notation in the transmission model. Transmission of the population ***q***^*B*^ with bottleneck *N* ^*T*^ results in the founder population ***q***^*F*^. The founder population grows under the influence of genetic drift, the effects of which are described by the effective population size *N* ^*G*^. Growth results in the population ***q***^*A*^. The populations ***q***^*B*^ and ***q***^*A*^ are observed, producing datasets represented by ***x***^*B*^ and ***x***^*A*^, which are used to reconstruct the original populations in terms of haplotypes. In order to calculate the variance of the reconstructed populations ***q***\*^*B*^ and ***q***\*^*A*^, datasets equivalent to ***x***^*B*^ and ***x***^*A*^, denoted ***x***\*^*B*^ and ***x***\*^*A*^ are generated and used to infer sets ***q***\*\*^*B*^ and ***q***\*\*^*A*^.

In generating the variance of our estimates, we use ***q***\*^*B*^ and ***q***\*^*A*^ to generate simulated observations, which we term ***x***\*^*B*^ and ***x***\*^*A*^. These in turn are used to generate a new round of estimates ***q***\*\*^*B*^ and ***q***\*\*^*A*^. In so far as ***q***\*\*^*B*^, ***q***\*\*^*A*^, ***q***\*^*B*^, *l* ^*l l*^ and ***q***\*^*A*^ are all known, they may be used to estimate the variances of ***q***\*^*B*^ and ***q***\*^*A*^.

### Haplotype reconstruction

We developed a maximum likelihood approach for haplotype reconstruction based upon existing technologies for processing short read data (24, 25, 27). We here assume that we have short-read data describing a viral population both before and after a transmission event. Before commencing haplotype reconstruction we performed three steps to pre-process the data using our software package SAMFIRE (25). Firstly, after alignment to the viral genome using BWA (28), the short read data were filtered, trimming reads to achieve a median PHRED score of at least 30, combining data from paired-end reads, and removing individual base calls with a PHRED score less than 30. Secondly, the filtered data were used to identify loci at which a polymorphism existed at significant frequency, this being defined using a cutoff of 2%. Thirdly, reads were processed to generate partial haplotypes, which describe the nucleotides present at each of the polymorphic loci in each read. Partial haplotype data were divided into distinct sets of reads, each describing alleles at a distinct set of loci in the viral genome. As an optional step, an estimate may be produced of the extent of noise present in sequence data, inferring a parameter, *C*, which describes the precision with which measurements of allele frequencies may be calculated via sequencing (25). A value of *C* = 1 here corresponds to a case in which reads are completely uninformative, while large values of *C* tend towards the binomial case in which each read accurately describes the allele present in a distinct viral genome, sampled in an unbiased manner from the population. A default value of *C* = 200 was used for our simulations.

We denote the sets of partial haplotype data collected before and after transmission as 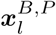 and 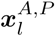 respectively, where *l* denotes the partial haplotype set. We now suppose that the viral population is comprised of a set of distinct haplotypes, denoted ***H***, which comprises *k* haplotypes, having the frequencies 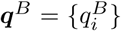 before transmission and 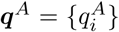 after transmission. These frequencies can be converted into partial haplotype frequencies by projection of the full haplotype space onto each lower-dimensional partial haplotype space by means of matrices *T*_*l*_. For example, given the full haplotypes before transmission {GA, TA, GC, TC} and a set of partial haplotypes {G-, T-}, we may write

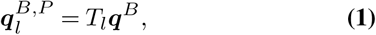

or more explicitly,

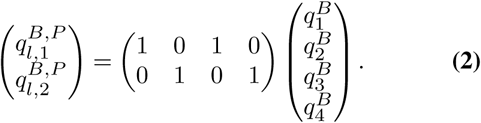

In the above instance we note that each partial haplotype can potentially be emitted from at least one of the haplotypes in ***H***. In order to generalise our model, we included in each set ***H*** a further haplotype ‘X’, describing the cloud of all potential viral haplotypes of the same length as those in ***H***, yet not already defined as being in ***H***. With this inclusion, we may say that any potential partial haplotype may be emitted from at least one of the haplotypes in ***H***, being emitted either from one of the defined haplotypes or from ‘X’.

In this way, we can construct a likelihood for any given set of haplotypes and frequencies, given the partial haplotype data. We write:

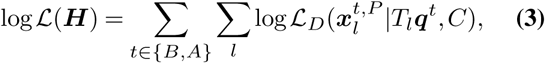

where ℒ_*D*_ denotes the Dirichlet multinomial likelihood

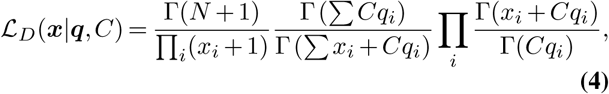

in which *N* = ∑_*i*_ *x*_*i*_.

A two-step optimisation was used to infer the optimal set of haplotypes and frequencies. To construct an initial set ***H***, a set of *k* ≥ 1 unique haplotypes were created in turn, to which was added the additional X haplotype. The frequencies of these haplotypes before and after transmission were then optimised under the constraint that the frequency of the X haplotype could not be greater than 0.01; this prevents the inference of trivial solutions to the model. We denote the inferred haplotype frequencies as ***q***\*^*B*^ and ***q***\*^*A*^. We note that the frequency of the X haplotype may be effectively zero; for the purposes of calculation a minimum frequency of *ϵ* = 10^−20^ was imposed.

Given our likelihood function, a series of changes were made to the set ***H***, optimising the frequencies each time to find the optimal haplotype reconstruction. Repeating this for increasing values of *k* gives a series of fits to the data; we used the Bayesian Information Criterion (BIC) to distinguish the most parsimonious explanation for the data:

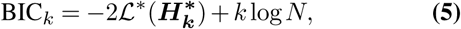

where 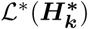 is the optimum likelihood value for the optimal set 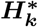 of *k* haplotypes, and *N* is the total number of observations in the dataset. Optimisation of the haplotype set was conducted for increasing values of *k* until a model with an additional haplotype produced an improvement of less than 10 units of BIC, representing a conservative cutoff point; a smaller required improvement would lead to the inference of a greater number of haplotypes. We note that the same haplotypes must explain both the samples collected before and after transmission; cases where haplotypes died out in transmission or were created *de novo* following the transmission event were inferred.

### Estimated error in reconstructed haplotype frequencies

For our compound method for bottleneck inference, we require an estimate of the variance in the inferred haplotype frequencies ***q***\*^*B*^ and ***q***\*^*A*^, so as to account for noise in these parameters when evaluating changes in the population. Variances were calculated by means of simulated data. Considering data collected before transmission, we used the frequencies ***q***\*^*B*^ to generate sets of partial haplotype data 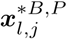, where *j* is used to index different sets. Each set provided an independent statistical replicate of the original data; having an identical number of sets of partial haplotypes, each spanning the same loci and containing the total number of samples. Each set was generated using a random Dirichlet multinomial sampling process with value *C* identical to the original. For each set of data, the haplotype reconstruction process was repeated, but with the haplotypes ***H*** constrained to those inferred for the original data. This process was repeated for 100 sets of data, generating the inferred haplotype frequencies 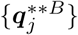. These values were used to calculate the diagonal elements of a covariance matrix var(***q***\*^*B*^) for ***q***\*^*B*^, given by:

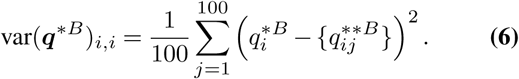

For simplicity, off-diagonal elements of this matrix were set to zero. An identical process was used to generate the matrix var(***q***\*^*A*^).

### Allele-frequency model of bottleneck inference

In generating Figure 1 we used a simple single-locus model of bottleneck inference. Given a set of independent allele frequencies 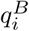 in the pre-transmission viral population, and their equivalent values 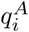 in the post-transmission population, we note that in the absence of selection, the mean value of 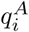 is given by 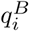, while the variance of 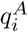, arising from genetic drift in a haploid system is given by

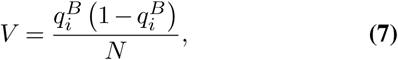

where *N* is the effective population size of the system (29).

To estimate the bottleneck size at transmission, we made the approximation that 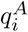 is normally distributed, then maximised the sum of the log likelihood values across allele frequencies

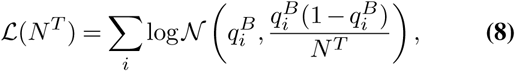

where *N*^*T*^ is the transmission bottleneck.

### Haplotype-based methods of bottleneck inference

Frequencies inferred from the haplotype reconstruction were used for the explicit and compound methods for calculating bottleneck size. As a first step in each method we removed haplotypes that were inferred to have been created *de novo* in the recipient following the transmission by removing haplotypes for which the pre-transmission frequency fell below a threshold frequency *δ*, set by default to 0.5%. Elements of the vectors ***q***\*^*B*^ and ***q***\*^*A*^ and the respective rows and columns of their covariance matrices were removed in this preliminary step.

In so far as we consider influenza transmission, we consider data from each viral segment independently, calculating first a likelihood of the bottleneck size given data from each segment, before combining the likelihoods across segments to estimate an overall maximum likelihood value for the transmission bottleneck.

### Compound method for bottleneck estimation

In the case of larger values of *N*^*T*^, an approach building upon that described in a previous publication (17) was applied. Briefly, we note that in a neutral transmission bottleneck, the expected composition of the population in the recipient is identical to that in the original host. The variance in this population is then a function of the size of the bottleneck and the extent of genetic drift during within-host growth, while in the case of inference, variation arising from the measurement of each population must also be considered.

Similarly to the approach outlined in an earlier work (17), we calculate a likelihood function with two components:

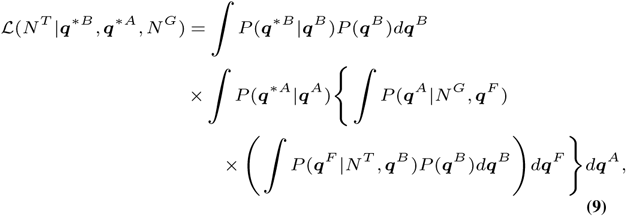

where the first integral corresponds to the initial observation of the system and the second encompass transmission (with the bottleneck *N*^*T*^), within-host growth (with drift described by the effective size *N*^*G*^) and post-transmission sampling. Each component of the likelihood is relatively simple to consider, as either a multinomial or Dirichlet-multinomial process, but the compound is difficult to evaluate. We note that, in cases where the frequency of a haplotype remains far from 0 or 1, and in particular as *N*^*T*^, becomes large, the likelihood can be increasingly well approximated in terms of a Gaussian distribution, with mean and variance calculated below.

Our solution makes use of the laws of total expectation and total variance. Given distributions *U* in *x* and *V* in *y*, the compound distribution *W* takes the form

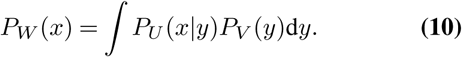

The mean and variance of *W* are then defined by

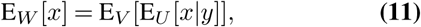

and

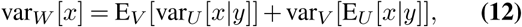

respectively.

For the pre-transmission component, the calculation of mean and variance are simple; our haplotype reconstruction process gives the estimate

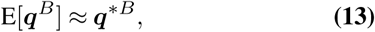

where the right-hand side is the output of the haplotype reconstruction, and

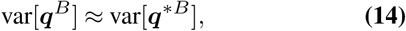

where the right-hand side was calculated using the generation of the datasets 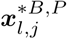 and the inferences of the frequencies {***q***\*\*^*B*^}_*j*_.

Moving on to the post-transmission component of the compound distribution in Equation 9, we can carry out the relevant marginalisations using the law of total expectation and the law of total variance.

Given that the dynamics governing transmission and within-host growth are assumed selectively neutral, the mean frequencies of the viral population are unchanged following transmission and growth. The mean term is therefore straightforward to calculate.

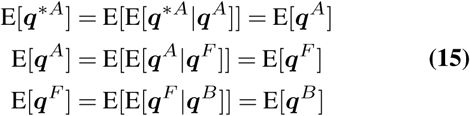

Thus

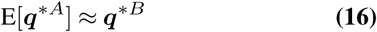

Calculation of the variance requires a little more effort. The transmission event can be modelled as a single multinomial draw with *N*^*T*^ number of trials. As a result, the variance of the founder population is given by

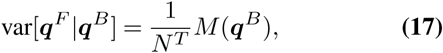

where *M* (***q***) = Diag(***q***) *−* ***qq***^†^.

We therefore obtain that

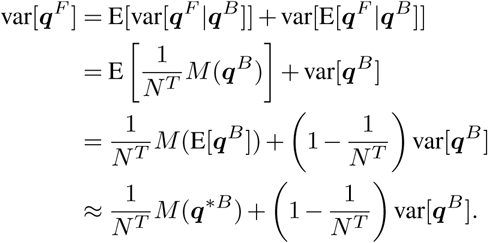

The within-host growth dynamics can be modelled as a multinomial draw of depth *N*^*G*^ = *gN*^*T*^ where *g* is the growth factor. From this we obtain the result that

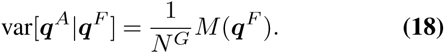

Marginalising over ***q***^*F*^ we obtain the variance

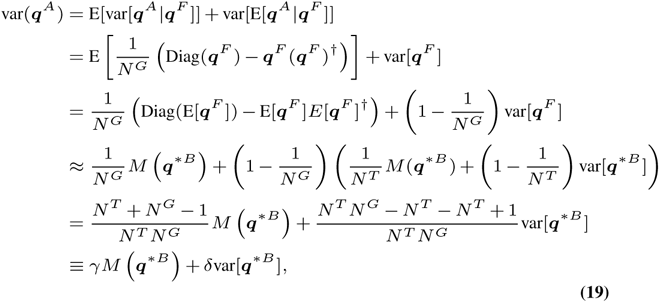

where we define 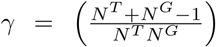and 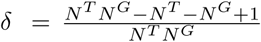.

Finally we have that

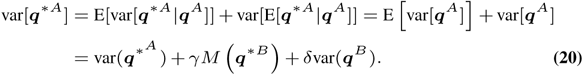

Together, Equations 15 and 20 define the mean and variance of a multivariate normal distribution representing the post-transmission component of the likelihood in Equation 9. Given our inferences for ***q***\*^*B*^ and ***q***\*^*A*^, we optimised the likelihood with respect to *N*^*T*^, generating a maximum likelihood estimate for the bottleneck size. We note that our approximation of the likelihood in terms of a multivariate normal distribution, works best where individual haplotype frequencies are not too close to zero or one, and where *N*^*T*^ is large. However, the approach allows for rapid calculation. In this sense we say that the compound method is optimised for large *N*^*T*^.

### Correction for the extinction of haplotypes in the compound method

Where a haplotype goes extinct in the transmission process, the likelihood function of the compound method can provide a poor estimate to the correct value. In this special case, relevant in our simulated data, we used a conditional distribution approach to make a correction to the likelihood.

In the above approximation we generated a multivariate normal distrbution for ***q***\*^*A*^:

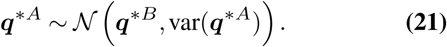

In this context, we split the vector ***q***\*^*A*^ into 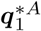 and 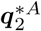, the latter containing all haplotypes post-transmission with a frequency lower than the threshold frequency *η*, which were considered to have died out during transmission, with the former containing the ‘surviving’ haplotypes. Rows and columns of the vectors and matrices were rearranged to put equation 21 into the form

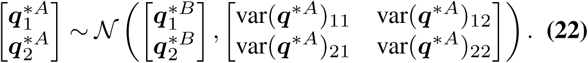

The frequencies of the components of the vectors were renormalised, such that 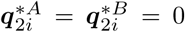, while 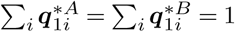.

We obtain the result that the conditional distribution of 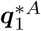 has the mean

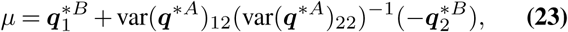

and covariance matrix

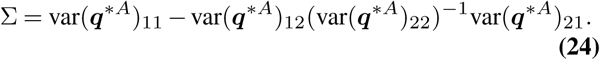

Using these parameters to define a Gaussian distribution, we calculated the likelihood of a bottleneck *N*^*T*^ given the data for the surviving haplotypes represented by 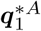.

To account for the haplotypes which became extinct during transmission, we made the assumption that these died out at the point of transmission to the founder population, the rapid growth of the founder population ensuring that no haplotypes went extinct through genetic drift, and viral sequencing of a large number of viral particles ensuring that no haplotypes were missed by the sequencing process. Under this assumption the likelihood of extinction is given by the simple binomial likelihood

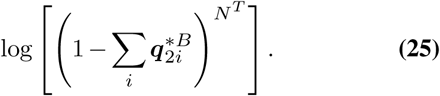

Summing the log likelihoods calculated for the surviving and the extinct haplotypes gave the total likelihood of the bottleneck size *N*^*T*^; the maximum likelihood value was identified via a simple optimisation process. To prevent nonsensical outcomes at very low bottleneck sizes, we further imposed the constraint that *N*^*T*^ could not be less than the number of haplotypes which survived transmission.

### Explicit method for bottleneck estimation

The explicit method uses the inferred haplotype frequencies for the population before transmission to reconstruct the space of possible outcomes in the recipient individual. Given our inferred haplotype frequencies 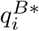, we assume that *N*^*T*^ viruses are transmitted. The probability that the founding viral population includes *n*_*i*_ copies of the haplotype *i*, where Σ_*i*_ *n*_*i*_ = *N*^*T*^, is given by

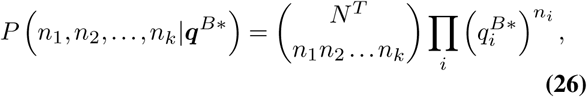

where the first term in the right-hand side of the equation is the multinomial coefficient.

For each possible outcome {*n*_*i*_} of this multinomial process, we obtained an inference of the haplotype composition 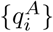 of the transmitted population given the relationship 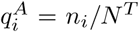 for each haplotype *i*. We then calculated the raw likelihood of observing the partial haplotype data collected post-transmission given this composition using the Dirichlet multinomial formulation described above, summing likelihoods over the possible outcomes of the initial transmission.

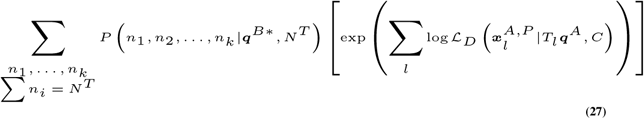

In this way we evaluate the likelihood of the bottleneck size *N*^*T*^ given the inferred pre-transmission haplotypes ***q***^*B*^ and the observed sequence data ***x***^*A*^; this is in contrast to the compound method, which is based on ***q***^*B*^ and ***q***^*A*^. We note that this approach neglects an explicit accounting for within-host growth of the population. Different assumptions about the dynamics of early viral infection can lead to changes in inferred bottleneck sizes (17); we are not confident that the biological reality of this phenomenon is well understood. Modifications to the the Dirichlet multinomial distribution could potentially be used in this context; increasing the variance of the likelihood function would soften the effect of small changes in the underlying population.

This approach has both the advantage and the disadvantage of explicitly representing the full set of all possible multinomial outcomes of transmission. While in this sense it remains close to the biological reality, it rapidly becomes computationally expensive as the number of haplotypes *k* increases and as *N*^*T*^ becomes large. For this reason we propose it as being optimal for small values of *N*^*T*^.

We note that, in our application to data from a transmission study presented here, the case in which a high bottleneck was inferred involved very limited diversity within viral segments; this facilitated the application of this method to consider larger bottleneck sizes.

### Generation of simulated data

Simulated data were generated using a simplified model of influenza transmission. Viruses were generated to have eight independent segments, of lengths equal to the segments of the A/H1N1 influenza virus. Each segment had five uniformly distributed polymorphic loci, making a theoretical total of 32 full haplotypes. Eight haplotypes were chosen from this set under the constraint that each of the five loci had to remain polymorphic. The frequencies of these haplotypes were then randomly generated under the constraint of a minimum haplotype frequency of 5%.

Each transmission event was modelled as a simple multinomial draw, selecting a number of viruses equal to the bottleneck size from the donor population. Within-host growth was then modelled as a second multinomial draw, conferring a 22-fold increase in the population size (30). Partial haplotype data were generated from simulated short reads of each viral segment. Short reads with lengths derived from the dataset of a recent influenza study (31) were generated (mean read length = 119.68, SD read length = 136.88, mean gap length = 61.96, SD gap length = 104.48, total read depth = 102825), these reads being used to calculate the number of reads spanning each set of consecutive polymorphisms in each segment. Given these numbers, partial haplotype observations were generated using a Dirichlet multinomial sampling process.

An inference of the transmission bottleneck was carried out independently using simulated data from each viral segment. These inferences were then combined, summing the log likelihoods across different segments to obtain an overall maximum likelihood estimate. Within our simulated data a small number of cases were identified in which the entire post-transmission population in a segment was inferred to comprise a haplotype that was not present above the cutoff frequency in the pre-transmission population, equivalent to a case where a haplotype arose *de novo* in the population and swept to fixation before data could be collected. In such cases, data for the segment in question were ignored, calculating the transmission bottleneck across the remaining segments.

### Processing of sequence data

Our method was applied to data from a recent study of influenza transmission among individuals in households (3). Data from transmission pairs identified in this study were aligned using the BWA software package (28) then filtered using SAMFIRE (25) to remove reads with a median PHRED score below 30, and to mask nucleotides with a PHRED score below this value. Polymorphic sites in coding regions of the virus were then called at an allele frequency cutoff of 2%, following which reads were divided into sets of partial haplotype data.

Data describing the within-host evolution of influenza were used to evaluate the extent of noise in the dataset. Where multiple samples were collected from a single host, trajectories were generated describing the change in each allele frequency over time. We have previously shown that an over-estimate of the extent of noise in sequence data can lead to substantial errors in the inference of a transmission bottleneck (17). Here, we used data from all single-locus trajectories to generate a provisional estimate of the extent of noise in the data. Trajectories which, under this estimate, evolved in a manner consistent with selective neutrality were then used to produce a final estimate of noise in the data; we inferred the parameter *C* = 660. Data from 43 putative transmission events were evaluated.

The estimate of an effective read depth for the case in which a very high bottleneck was inferred was conducted using SAMFIRE based upon allele frequency data, and using a cutoff frequency for minority alleles of 2%.

## Supplementary Note 1: Availability of code

Code and data used or generated during this project is available from https://github.com/cjri/VeTrans.

## Supplementary Note 2: Acknowledgments

This work was supported by a Sir Henry Dale Fellowship, jointly funded by the Wellcome Trust (wellcome.ac.uk) and the Royal Society (royalsociety.org) with grant number 101239/Z/13/Z. CKL was funded by a Wellcome Trust Studentship with grant number 105365/Z/14/Z. CJRI acknowledges a visiting fellowship from the University of Helsinki. DBW was funded by a Simons Investigator award from the Simons Foundation with grant number 508600.

